# Inferring the direction of rhythmic neural transmission via inter-regional phase-amplitude coupling (ir-PAC)

**DOI:** 10.1101/465484

**Authors:** Bijurika Nandi, Peter Swiatek, Bernat Kocsis, Mingzhou Ding

## Abstract

Phase-amplitude coupling (PAC) estimates the statistical dependence between the phase of a low-frequency and the amplitude of a high-frequency component of local field potentials (LFP). Characterizing the relationship between nested oscillations in LFPs, PAC has become a powerful tool for understanding neural dynamics in both animals and humans. In this work, we introduce a new application for this measure to two LFPs to infer the direction and strength of rhythmic neural transmission between distinct networks. Based on recently accumulating evidence that transmembrane currents related to action potentials contribute a broad-band component to LFP in the high-gamma band, we hypothesized that PAC calculated between high-gamma in one LFP and low-frequency oscillations in another would relate the output (spiking) of one area to the input (soma/dendritic postsynaptic potentials) of the other. We tested this hypothesis on theta-band long range communications between hippocampus and prefrontal cortex (PFC) and theta-band short range communications between different regions within the hippocampus. The results were interpreted within the known anatomical connections predicting hippocampus→PFC and DG→CA3→CA1, i.e., theta transmission is unidirectional in both cases: from hippocampus to PFC and along the tri-synaptic pathway within hippocampus. We found that (1) hippocampal high-gamma amplitude was significantly coupled to theta phase in PFC, but not vice versa; (2) similarly, high-gamma amplitude in DG was significantly coupled to CA1 theta phase, but not vice versa, and (3) the DG high-gamma-CA1 theta PAC was significantly correlated with DG→CA1 Granger causality, a well-established analytical measure of directional neural transmission. These results support the hypothesis that inter-regional PAC (ir-PAC) can be used to relate the output of a “driver” network (i.e., high gamma) to the input of a “receiver” network (i.e., theta) and thereby establish the direction and strength of rhythmic neural transmission.

## Introduction

Neural information processing depends on interactions between different groups of neurons. Being able to assess the patterns of neuronal interactions is essential for a better understanding of the cooperative nature of neuronal computation. Neural interactions are directional and involve different spatial and temporal scales. Cross correlation functions in time domain and coherence functions in spectral domain are often computed on neural signals recorded from distinct networks. Although these measures provide useful information regarding functional connectivity, they are technically symmetric in that when signal A is coherent with signal B, signal B is equally coherent with signal A, thus failing to contribute towards providing information on the direction of neural transmission. Cross-frequency coupling (CFC) is a measure increasingly used to quantify the relation between neuronal activities in different frequency bands within a single local field potential (LFP) recording. When applied to pairs of LFP signals recorded in distinct structures 1, however, CFC is inherently asymmetric; it can associate one feature in one LFP with a different feature in the other LFP signal. Based on this property of CFC, we advance here a new interpretation of this quantity, and suggest that it can be used to infer the direction and strength of rhythmic neural transmission between distinct brain networks.

Among the various CFC measures, phase-amplitude coupling (PAC), which estimates the statistical dependence between the phase of a low frequency oscillation and the amplitude of a high frequency oscillation, is the most commonly used. When applied to a single LFP recording, PAC results are often interpreted as a measure of how low-frequency fluctuations in membrane potential influence higher-frequency oscillations, namely, greater activity of excitatory and inhibitory neuron populations generates stronger postsynaptic potentials at high frequencies. We note that the implicit assumption that PAC represents the influence of low-frequency phase on high-frequency amplitude does not necessarily apply in all situations; the opposite influence may take place in a two-LFP scenario in which high-frequency activity of one LFP, taken to reflect population spiking, causes low-frequency oscillations in the membrane potential of the other LFP. In particular, when the pair of LFP signals are recorded in distant structures where establishing inter-regional synchrony requires the transmission of spike activity, the second interpretation is likely more accurate.

Owing to the ambiguity of the origin of LFP 2, the interpretation of CFC measures is not always single-faceted. LFP is mainly generated by postsynaptic potentials but may also contain wide-band signals resulting from spikes and spike afterpotentials 3, 4. Neuronal spikes were shown to correlate with gamma-band power in cognitive tasks 5, 6. Recently, accumulating evidence indicates that LFP contains broad-band high-gamma (> 60 Hz) components generated by fast transient transmembrane currents related to action potentials 7, 8. Buzsaki and Wang (2012) even noted that, “although spike contamination can be a nuisance, by using proper analytical methods, spike power can be exploited as a proxy for the assessment of neuronal outputs even in recordings of LFPs.”

The foregoing laid the foundation for the proposed extension of single LFP PAC to two LFPs in which we use the fast component in the high-gamma range as an indicator of spike activity representing the output of one network and test whether it correlates with low-frequency oscillations in structures representing downstream targets of this network. We hypothesized that for two neural networks with a driver-receiver relationship, PAC calculated from LFP recordings from the two networks, referred to inter-regional PAC (ir-PAC) here, will be significant in one direction only, i.e., between high-gamma amplitude of the “driver”, representing rhythmically modulated spike trains, and the phase of the low-frequency oscillation in the “receiver” network, representing the resultant postsynaptic potentials (see Fig. 1 for a schematic). We tested this hypothesis on different datasets involving simultaneous LFP recordings from PFC and different regions of the hippocampus where directionality is well-established by abundant anatomical and physiological evidence. We predicted that hippocampal high-gamma amplitude was significantly coupled to theta phase in prefrontal cortex (PFC), but not vice versa, and high-gamma amplitude in dentate gyrus (DG) was significantly coupled to CA1 theta phase, but not vice versa, in agreement with known anatomical connections predicting unidirectional theta transmission from hippocampus to PFC 9-11 and along the tri-synaptic pathway within the hippocampus, but not back. For data where the appropriate experimental conditions were met 12, the direction of DG→CA1 theta drive was also verified by Granger causality (GC) 13, a well-established analytical measure of directional neural transmission (Ding et al., 2006). For further support, local PAC (l-PAC) was also calculated between theta frequency and high-gamma amplitude in each of the recorded two structures, and we predicted that l-PAC was significant in the “driver” circuit, indicating that rhythmic membrane potential oscillations drive rhythmically synchronized spike train output, but significant l-PAC did not necessarily appear in the “receiver” circuit, depending on whether the rhythmic postsynaptic potentials was subthreshold or suprathreshold.

**Figure 1.**
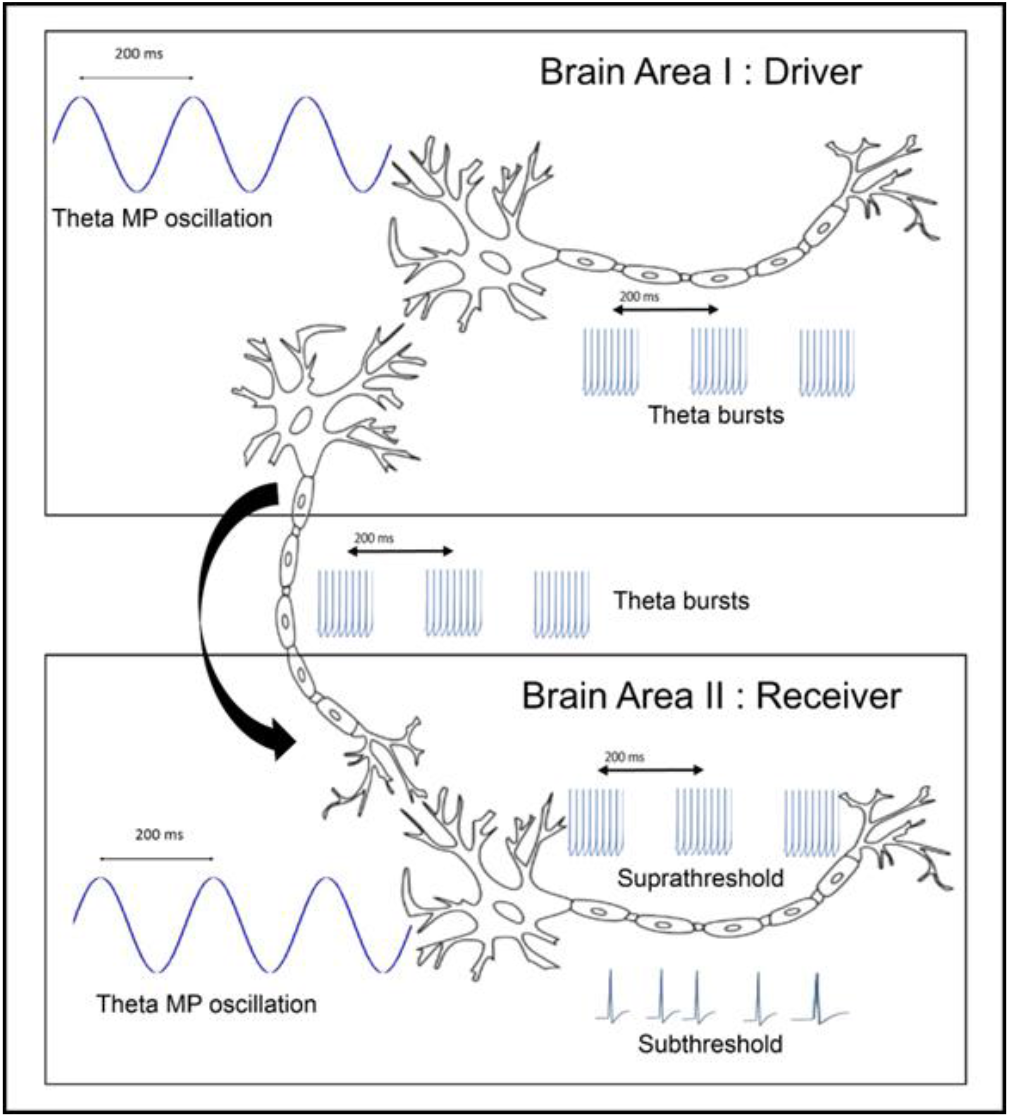
The origin of electrical signals generating significant ir-PAC. Slow rhythmic (e.g. theta) fluctuation of the membrane potential of principal neurons (theta MP oscillations) give rise to rhythmic firing of neuronal ensembles (theta bursts), generating the low and high-frequency components of LFP recorded within the same brain area and leading to significant local PAC. Neurons of this region (Brain Area I) will also send rhythmic bursts through their fiber output pathways to target areas (Brain Area II) where they will elicit rhythmic synaptic potentials on the input segment of local neurons. Thus, the signal representing the output segment of neurons in the “driver” (gamma related to firing, Brain Area I) will be statistically related (i.e., strong ir-PAC) to the slow LFP “receiver” (membrane potential oscillations in Brain Area II). Depending on firing threshold of individual neurons in the receiver, local PAC may or may not be generated in this area.

## Results

Our theoretical consideration behind ir-PAC is illustrated in Figure 1 for a pair of networks forming a driver-receiver relationship. In Brain Area I, action potentials bursting at the theta frequency contribute to the high gamma component of the LFP recordings in the driver network. These actions potentials were transmitted from Brain Area I to Brain Area II. The membrane potential (MP) responses to the action potential input contribute to the theta component within the LFP recordings in the receiver network. The proposed ir-PAC assesses this relationship. Depending on whether the MP oscillations are suprathreshold or subthreshold, the local PAC or l-PAC may or may not be statistically significant; in other words, the action potentials in the receiver network may or may not be theta rhythmic.

We tested the possible utility of using ir-PAC to infer direction of rhythmic neural transmission on data recorded from brain networks exhibiting theta rhythm. See Figure 2 for illustrations of the three datasets. Theta, known to originate in the hippocampus and transmit from the hippocampus to other parts of the brain in a behavior- and state-dependent manner ^14, 15^, has been intensively studied, and the fund of knowledge in the extant literature provides the basis for evaluating the performance of the proposed method.

**Figure 2.**
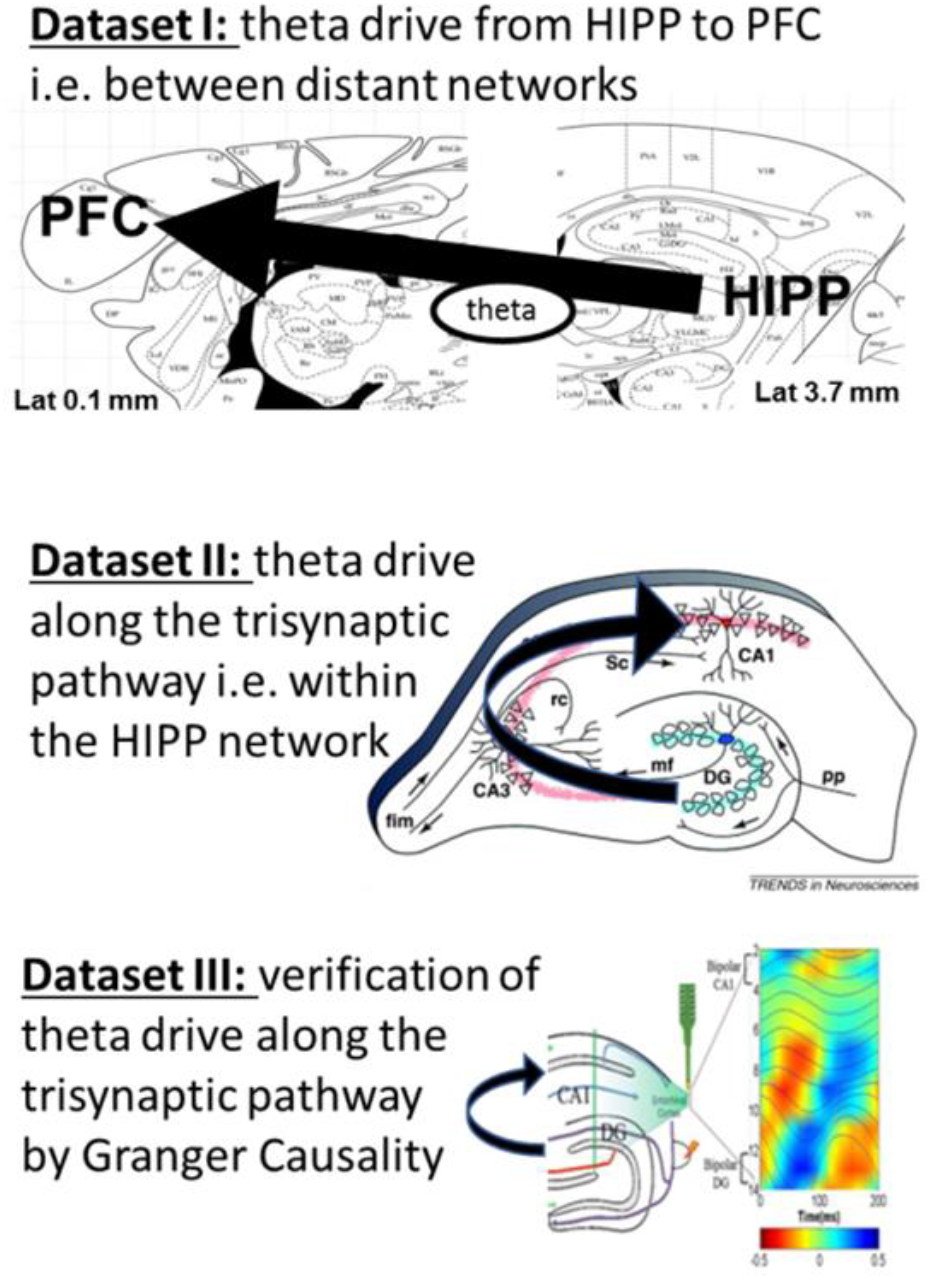
Schematics of analysis used in this study to test the hypothesis that ir-PAC across distinct areas can be used to estimate directionality of neural transmission between them. We used 3 datasets, with hippocampal theta drive spreading to the PFC (distant network) carried by rhythmic firing in the HIPP →PFC pathway (Dataset I) or within the hippocampus along the classic trisynaptic pathway from the electrode placed below the fissure to the electrode above the CA1 pyramidal layer (Datasets II and III). Datasets I and II included unipolar LFP recordings in two locations during theta states (waking exploration and REM sleep) in freely moving rats. Dataset III compared ir-PAC and GC directions using theta LFP evoked by RPO stimulation and recorded by a 16-channel silicon probe across CA1 and DG in urethane-anesthetized rats. Theta dipoles in DG and CA1 were identified using perforant path stimulation, and PAC and GC was calculated between bipolar recordings, based on CSD analysis ^12^.

Amplitude-phase pairing in PAC calculation, i.e., gamma amplitude in the driver network with theta phase in the receiver network or vice versa, was calculated to examine whether it corresponded to known anatomical connections, and matched the direction identified by Granger causality when the dataset was appropriate for a GC analysis. In the first two datasets, we used unipolar LFP recordings in hippocampus and PFC (Dataset I) and in DG and CA1 (Dataset II) as examples of theta coupling between distant structures and between local networks within the hippocampus, respectively. These two datasets were recorded in freely moving rats during behavioral theta states of awake exploration and REM sleep. In the third dataset (Dataset III), we used 16-channel laminar LFP recordings across hippocampal layers, extending from above the CA1 pyramidal layer to the DG below the hippocampal fissure. Bipolar LFP derivations representing theta dipoles elicited by brainstem stimulation in urethane anesthetized rats were made to allow precise measurement of GC ^12^; GC was used for verification of the direction and strength of theta drive within the hippocampus estimated by ir-PAC (Fig. 2).

In Dataset I, analysis of LFP recordings in the hippocampus and PFC during natural theta states tested the possible theta drive between two distant structures along the anatomically demonstrated unidirectional hippocampo-prefrontal pathway ^9-11^. There was strong local theta versus high-gamma PAC in hippocampal recordings of all experiments, both during awake exploration and REM sleep (Fig. 3), corresponding to known synchronized theta fluctuations in membrane potential and in firing activity ^16, 17^ during these states. l-PAC reveals the connection between the two frequencies but does not reveal causality, i.e., whether membrane potential oscillations modulate spike firing or rhythmic action potential bursts are generating rhythmic synaptic membrane potentials. l-PAC was low in the PFC (Fig. 3B), where no significant coupling was found in the REM segments of 6 recordings, even though it was significant during waking occasionally (in 3 out of 6 recordings) in individual experiments (see, e.g., Experiment M74 in Fig. 3A), possibly depending on behavior which was not controlled in these experiments. Ir-PAC revealed significant coupling (p<0.05) across hippocampal and PFC LFPs during theta states of both waking and REM sleep. Specifically, ir-PAC was high (significant coupling in 4 out of 6 recordings) between hippocampal high-gamma amplitude and PFC theta phase (ir-PAC=3.2±1.5 and 4.4±1.9 in awake and REM, respectively) but was not significant when calculated using high-gamma amplitude in PFC and theta phase in hippocampal recording (ir-PAC=1.6±0.7 and 1.6±0.2 in awake and REM, respectively), indicating that the first pairing showed the significant HIPP→PFC theta drive, as predicted; this finding was in agreement with known physiology of theta rhythm, namely, it is primarily generated in the hippocampus, and propagated to PFC via HIPP to PFC connections.

**Figure 3.**
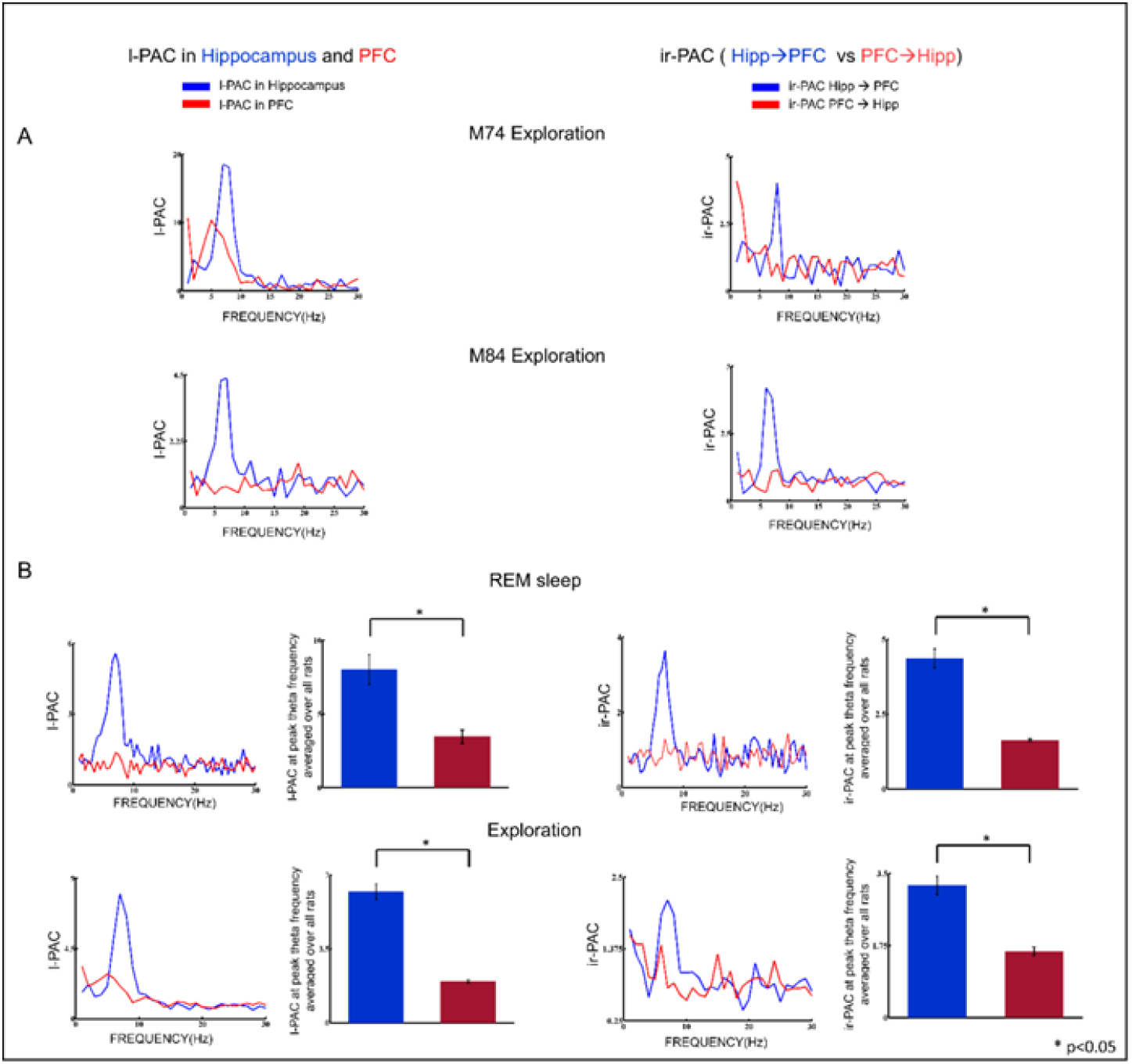
Theta-gamma PAC in two distant structures, hippocampus and PFC. *Left:* Local PAC calculated using low frequency phase (0-30Hz) and gamma (65-90 Hz) amplitude of the same signal in hippocampus (HIPP) and prefrontal cortex (PFC). *Right:* ir-PAC calculated using pairs of low frequency phase and gamma amplitude from signals recorded in different locations. **A**. Two examples showing strong local theta-gamma PAC in hippocampus and weaker (expt. M74) or no theta PAC (expt. M84) in PFC during wake exploration. Note significant ir-PAC calculated across areas, i.e., using hippocampal gamma and PFC theta, in both examples, but no significant ir-PAC between PFC gamma and hippocampal theta. **B**. Group averages (n=13) of local PAC and ir-PAC across hippocampus and PFC during REM sleep (*upper*) and exploration (*lower*). Note strong local theta PAC in hippocampus and no theta PAC in PFC. Theta carried from hippocampus to PFC (Hipp →PFC) is expressed in significant ir-PAC of PFC theta by hippocampal gamma.

Next, we analyzed interregional theta versus high-gamma PAC across different subnetworks within the hippocampus in freely moving rats in natural theta states of exploration and REM sleep (Dataset II) and in rats in which theta was elicited by brainstem stimulation under urethane anesthesia (Dataset III). In both datasets, the amplitude of gamma oscillations in DG was significantly coupled to the phase of theta oscillations in CA1, but not vice versa, implying unidirectional synaptic transmission from DG to CA1 and no theta transmission in the opposite direction, i.e., from CA1 to DG (Fig. 4). Thus, in both freely moving and anesthetized rats, significant ir-PAC (n=9 for Dataset II and n=2 in Dataset III) existed across areas between DG high-gamma amplitude and CA1 theta phase (ir-PAC=3.6±1.6 and 1.6±0.8 in Datasets II and III, respectively), but not vice versa (ir-PAC=1.5±0.4 and 1.3±0.4). The direction identified by ir-PAC corresponded to transmission along the unidirectional intrahippocampal trisynaptic DG→CA3→CA1 pathway, and lack of anatomical projection from CA1 to DG ^18-21^. It is noteworthy that the results were similar in the two datasets (Dataset II and III), even though PAC in Dataset II was calculated among unipolar hippocampal LFP recordings obtained with two electrodes placed in or above the CA1 pyramidal layer and below the hippocampal fissure in the DG respectively, whereas in Dataset III, LFPs in the two major theta dipoles were used utilizing bipolar recordings derived from 16-channel laminar LFPs. Also, in both Datasets II and III, the amplitude of gamma oscillations in DG was significantly coupled locally to the phase of theta oscillations of DG itself (l-PAC=2.9±1.5 and 1.8±1 in Datasets II and III, respectively), whereas l-PAC was not significant in CA1 (l-PAC=1.8±0.5 and 1.3±0.3 in Datasets II and III, respectively) (Fig. 4A).

**Figure 4.**
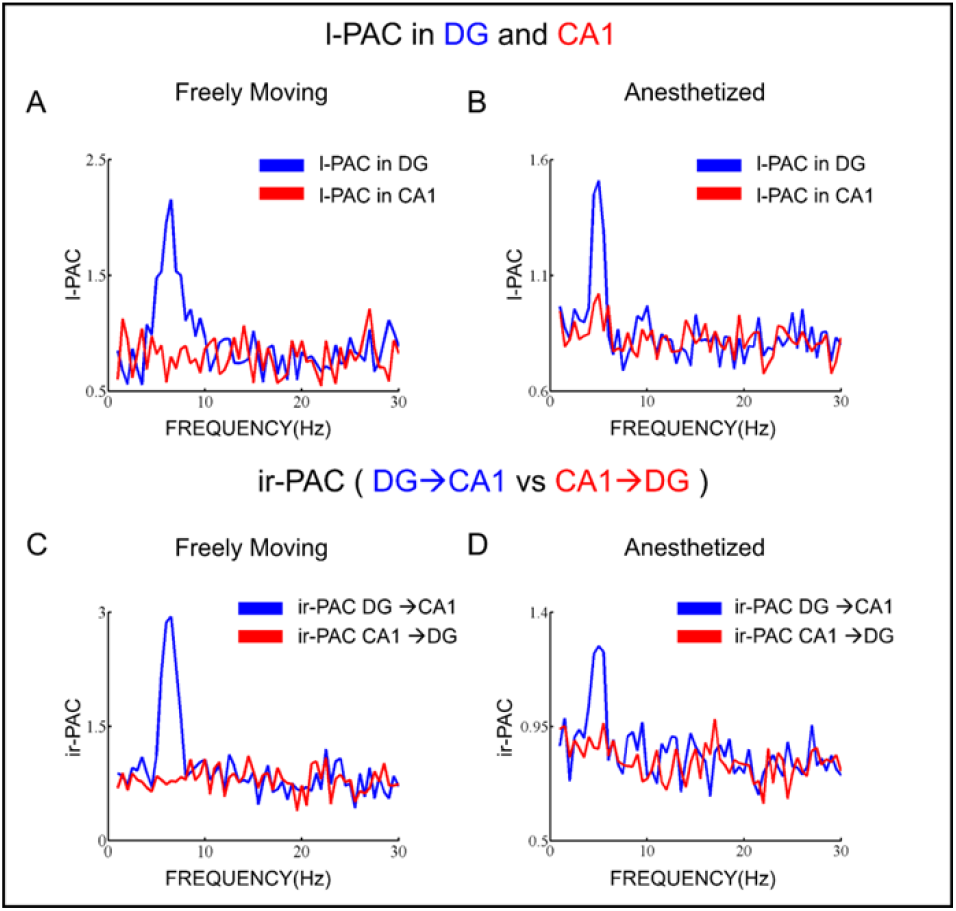
Theta-gamma PAC within the hippocampal areas of DG and CA1. Local PAC between gamma amplitude and theta phase in the frequency range of 1-30 Hz calculated using the same LFP signal in DG (*Blue*) and CA1 (*Red*), averaged across experiments with unipolar recordings of two LFP signals in freely moving rats in natural theta states *(n = 13)* (**A**) and experiments with bipolar recordings in the two major theta dipoles (derived from 16-channel LFP recordings across hippocampal layers of CA1 and DG) activated by brainstem stimulation under urethane anesthesia *(n = 9)* (**B)**. ir-PAC between DG gamma (65-85 HZ) amplitude and CA1 phase frequency (*blue*) and vice versa (i.e. between CA1 gamma and DG theta – *red*) in the range of 1-30 Hz; group averages in freely moving *(n = 13)* (**C**) and urethane-anesthetized rats *(n = 9)* (**D**).

In the foregoing analysis, we specifically focused on the upper gamma band (>60Hz) as a proxy of spike activity ^7, 8^. Over the entire gamma range of 30-100 Hz, theta-gamma PAC was high in the high-gamma band with peak in the 65-85 Hz range, considerably exceeding theta-gamma PAC in the low gamma range (<55 Hz) (Fig. 5). It is known in contrast, that membrane potential oscillations generated by multiple mechanisms in the gamma range involving different types of interneurons and serving different memory functions appear in narrower gamma bands distributed however both around 40 Hz and at higher frequencies ^2, 22^.

**Figure 5.**
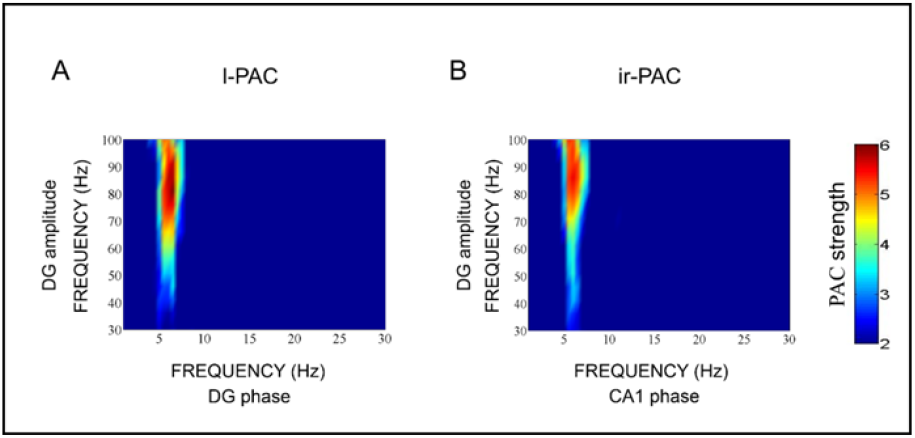
Role of wide-band gamma components in local and ir-PAC. Example of theta-gamma local PAC in DG (**A**) and ir-PAC (DG→CA1) (**B**) in a freely moving rat. Comodulograms were calculated between gamma amplitude (30-100 Hz) and theta phase (1 – 30 Hz). Note that both PAC measurements are limited to narrow theta band (5-7 Hz) but are distributed over broad gamma band (>60 Hz).

Existence of significant theta-gamma PAC was also strongly influenced by the strength of theta oscillations. To show this, we divided individual LFP recordings in Dataset II into two groups: (1) which had significant ir-PAC between DG gamma amplitude-CA1 theta phase (n=9) and (2) which did not show any significant PAC, either across areas or local in the driver network DG (n = 4). Average theta power in recordings that showed significant ir-PAC had higher power in both DG and CA1 than in the group showing no significant ir-PAC (Fig. 6).

**Figure 6.**
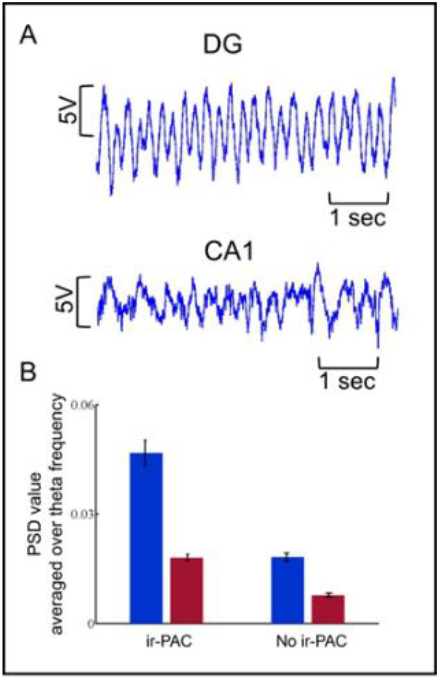
Relationship between theta power in DG and CA1 and theta-gamma ir-PAC across DG and CA1. **A**. Sample LFP traces (6 s) recorded in DG and CA1. **B.** Average DG and CA1 power in experiments with significant ir-PAC (n=9) and not significant (n=4) ir-PAC between DG gamma amplitude and CA1 theta phase.

For Dataset III, owing to the electrode configuration, bipolar derivations can be made for the theta dipoles, which enables the computation of Granger causality^13^. GC has been shown to be an effective method of evaluating causal relationships between neuronal ensembles ^12^. Thus, to verify the direction and strength identified by ir-PAC, we also calculated GC spectra of the same bipolar LFP recordings ^23, 24^ and averaged across all experiments under Dataset III (Fig. 2), following the procedures described earlier ^12^. GC revealed unidirectional information flow from DG to CA1 at theta frequency (~5 Hz in Fig. 7B) (GC= 0.94±0.52), whereas GC in the opposite direction showed no significant causality (GC value = 0.02±0.02). The pattern and the frequencies of theta corresponded in GC and ir-PAC measurements (Fig. 7A, B). Furthermore, calculated from the same LFP recordings, the magnitude of DG high-gamma amplitude and CA1 theta phase ir-PAC positively correlated with the magnitude of DG→CA1 GC (Fig. 7C) with a correlation coefficient of 0.9012 (p=0.0009). In contrast, there was no significant correlation (r = 0.4742, p=0.1972) between the magnitude of CA1→DG GC and ir-PAC calculated for CA1 gamma amplitude and DG theta phase (Fig. 7D). The comparison between the new technique, ir-PAC, and the well-established analytical measure of directionality, GC, was enabled by the significant variability of DG→CA1 direction measurements in individual recordings within Dataset III (Fig 8). ir-PAC was applied to unipolar wire recordings referenced to common reference in freely moving animals (Dataset II), whereas GC was applied to bipolar LFP recordings derived from 16-channel silicon probe across CA1 and DG in urethane-anesthetized rats (Dataset III). The first represents the main target of this application, the second represents the model allowing the most reliable GC estimations ^12^. GC appeared more reliable revealing significant DG→CA1 direction in all 9 recordings (Fig. 8B) whereas ir-PAC lead to significant estimate in 69% of (9 of 13, Fig 8A) recordings.

**Figure 7:**
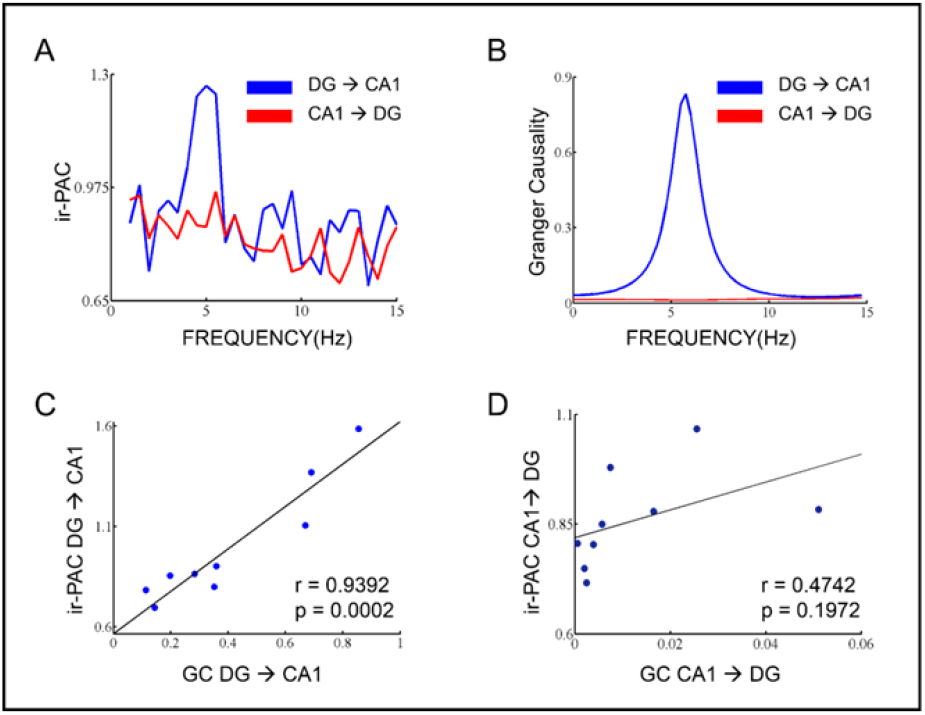
Comparison of directionality measures using ir-PAC and GC. **A.**Average ir-PAC strength between DG gamma amplitude and CA1 phase frequency range of 1-30 Hz and vice versa, averaged across all rats (n = 9). **B.** Granger Causality spectra between DG and CA1 averaged across all rats (n = 9). **C.** Correlation between GC (DG → CA1) values and ir-PAC (DG gamma amplitude - CA1 theta phase) values of all rats. **D.** Correlation between GC (CA1 → DG) values and ir-PAC (CA1 gamma amplitude - DG theta phase) values of all rats.

**Figure 8.**
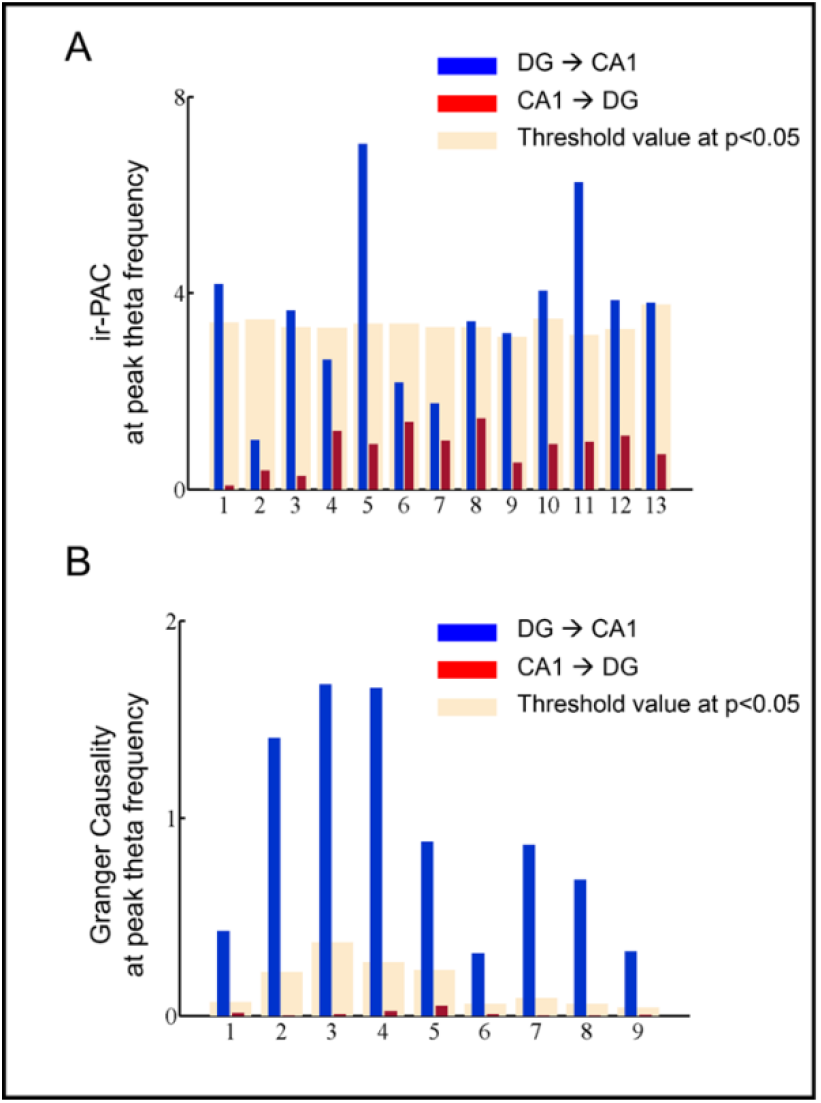
Comparison of estimates of the direction of within-hippocampal theta drive using ir-PAC of monopolar wire electrode recordings with GC calculated on bipolar recordings. **A.** Bar plots showing ir-PAC between DG high-gamma amplitude and CA1 theta phase (blue) for 13 unipolar wire recordings (Dataset II) as well as ir-PAC between CA1 high-gamma amplitude and DG theta phase (red); significance threshold (p<0.05) was indicated for each recording (cream). **B.** Theta-band DG →CA1 GC (blue) and CA1 →DG GC (red) for the 9 bipolar recording (Dataset III) recordings; significance threshold value at p<0.05 (cream).

## Discussion

In this study we tested the idea that ir-PAC, namely, the high gamma amplitude of one LFP and the theta phase of another LFP, can be used to infer the direction and strength of rhythmic neural transmission. Specifically, we found significant ir-PAC between high-gamma amplitude of LFP signals recorded in networks known to generate theta oscillations and theta phase of LFPs recorded in downstream structures receiving input from these networks through well-established non-reciprocal anatomical projections. When theta and high-gamma components of the LFP signals were switched, PAC showed no significant relationship, indicating that theta phase of the driver circuits and high-gamma amplitude of the receiver circuits are not coupled. These findings suggest that PAC can be used to infer direction and magnitude of rhythmic drive between distinct brain networks.

After formulating the idea conceptually, we proceeded to test it in two networks. The first network was the hippocampus-PFC network, a cortical network of two spatially segregated structures in which PFC neuronal firing and LFP were known to be modulated by hippocampal theta in a behavioral and task dependent manner ^15, 25-28^. Although theta rhythm can be transmitted to PFC and hippocampus from common sources or through a reciprocal connection through the thalamus, the dominant theta drive is carried by the non-reciprocal hippocampo-prefrontal pathway ^29^. In the second network, the DG and CA1 network within the hippocampus, theta is known to be generated in both recorded structures, by local mechanisms as well as common inputs, primarily from the medial septum and entorhinal cortex ^16-18^. Communication between DG and CA1 is unidirectional, however, through the classic trisynaptic pathway, thus indicating that there should be significant ir-PAC between DG high-gamma amplitude and CA1 theta-phase, but not the other way around, namely. This was also verified with GC, a well-established analytical measure of directionality which pointed in the same direction, i.e., DG→CA1, and showed numerical correlation with ir-PAC values (Fig. 7C).

The results are consistent with recently accumulating evidence that LFP recordings contain high-gamma components due to action potential transients in the underlying structure ^4, 7, 8^. In fact, postsynaptic potentials of true oscillations remain hard to separate from spike “contamination” ^7, 30-32^. Most spectral analysis techniques can reveal multiple oscillations as spectral peaks riding on top of the characteristic background 1/f decay in power spectra, but are inadequate to handle transients best described as delta function or a Gaussian with small sigma contributing to LFP spectra in a wide frequency range. Using matching pursuit, however, a technique that allows decomposition of LFP into oscillatory (narrow-band) and transient (broadband) components, Ray and Mansell (2011) demonstrated such LFP transients associated with spikes in specifically designed experiments in which gamma oscillations and spiking were anti-correlated in monkey visual cortex and could be controlled and thus be separated by changing the stimulation parameters. They also showed that although sharp transients most likely had power over the entire spectrum these could only be observed over ~50 Hz, due to masking at lower frequencies by the 1/f character of LFP spectra ^7^. The 50-60 Hz cut off frequency was further supported by detailed analysis of firing rate and band power in different behaviors showing positive correlation in a wide gamma range of 50-180 Hz with peaks between 60-90 Hz (^8^, their Fig.2). PAC between theta and broadband higher gamma followed a similar pattern in the present study (i.e., significant PAC broadly distributed above 55 Hz; Fig. 5). The 65-85 Hz frequency range chosen in our study for PAC calculation proved suitable to verify the hypothesis both for l-PAC within the same LFP recording and ir-PAC between LFPs in distinct structures, with the latter being exploited for directional information (Figs. 3, 4).

Although PAC calculated from one signal does not give directional information, it is tacitly assumed in most studies (see however ^33, 34^ for examples showing the opposite) that low-frequency oscillations modulate high-frequency oscillations, not vice versa. In contrast, in the present application of PAC between two LFPs, we found that, it was the high-gamma amplitude of one signal which was controlling theta phase of the other. Assuming that high-gamma reflect population spiking, therefore the output activity of a neuronal ensemble, and that low frequency oscillations (e.g., theta) reflect dendritic processing, therefore the input activity of a neuronal ensemble, we conclude that ir-PAC, instead of testing the relationship between a pair of genuine network oscillations, connects the output of a driver and the input of a receiver network. Thus, periodic increase in spike activity at the theta frequency in the driver network will result in theta bursts in hippocampo-fugal pathways which elicit theta rhythmic postsynaptic membrane potential response from neurons in the target receiver network. This interpretation is further supported by the finding that ir-PAC can only be measured effectively when slow frequency oscillations (e.g., theta) in the driver are strong, i.e., over the threshold of generating enough action potentials on the output. Our results (Fig. 6) show that datasets with low theta power in DG and CA1, did not show significant coupling in any direction, across or within areas.

The interpretation of two-LFP PAC measurements supported by the present findings suggests that ir-PAC may serve as a suitable tool to infer directional information for low-frequency rhythmic drive between distinct networks. Similar to other widely used techniques analyzing inter-regional communication, e.g. coherence, its effectiveness however is strongly influenced by the exact anatomical wiring defining the recorded LFP ^2^. In the laminar hippocampal datasets for example, which allow precise layer-specific localization of recordings (e.g., including the layer where CA3 theta input arrives in CA1 ^17, 18, 35-37^) and using bipolar constructs permitting the most reliable GC estimations ^12^, parallel ir-PAC and GC analysis showed a highly significant positive correlation between the two variables (Fig. 7), suggesting that ir-PAC reveals not only the direction but also the strength of rhythmic neural transmission. In the widely used experimental preparation in which hippocampal LFP is monitored in less stringent conditions, i.e. using monopolar CA1 and DG wire recordings, ir-PAC still lead to significant estimate in ~70% of recordings and pointed in the correct direction in all remaining experiments (Fig 8A), indicating that even when ir-PAC values failed to pass statistical threshold (thus deemed statistically not significant), the values may still be used to hint at the direction and strength of rhythmic transmission.

Theta-gamma PAC has become a widely used, powerful tool for understanding neural dynamics in both human and animal research. Gamma power in rat was found to be modulated by ongoing theta oscillations in rat hippocampus more than two decades ago ^38^. More recently, PAC was tested in a range of cognitive tasks; for example, theta-gamma phase-phase coupling was found during maze exploration and REM sleep ^39^, theta phase-gamma amplitude coupling strength increased during learning in rat hippocampus, thus contributing to memory processing ^40^, nesting of gamma frequency oscillations within slow theta frequency in rat hippocampus indicated routing of information as a major function of gamma frequency in different bands ^22^, high gamma power was found to be modulated by theta phase in human neocortex ^41^, simultaneous maintenance of multiple items in working memory was observed to be accompanied by PAC in human hippocampus ^42^, high frequency oscillation modulated by the type of low frequency oscillation is task dependent, as observed in human intracranial recording ^43^. The interpretation proposed for local PAC, l-PAC, may shed further light on these studies in terms of low-frequency oscillatory activities in a neural network and the relation with the output activities from the network. Similarly, the directional information obtained from ir-PAC may add a further dimension to the interpretation of the few prior studies in which PAC was calculated across regions and showed results equivalent to this study, i.e., significant PAC between the amplitude of high-frequency hippocampal LFP and theta phase of extrahippocampal structures, but not vice versa, including PFC (^1^; their Fig. 6C) and striatum (^44^; their Fig. 5A).

Two further observations are in order. First, although ir-PAC only gives information of propagation of the low-frequency oscillation, it has important impact on coupling of genuine gamma oscillations in distinct networks. Gamma oscillations are difficult to synchronize over longer distances because of their low power (compared with delta, theta, alfa, beta, - cf. 1/f spectral background) and uncertainties and non-uniformity of conduction times which have progressively higher impact as the oscillation cycle is getting shorter and shorter ^45^ (Buzsaki). Thus, gamma oscillations are considered local, i.e., generated by local networks in different cortical structures. Their synchrony is achieved by synchronizing low-frequency rhythms effortlessly carried over long distances entraining neuronal firing and establishing synchronized gamma oscillations on both ends of the pathway. Hippocampo-cortical communication is based for example on theta synchrony flexibly established between cooperating networks in a behavior and task-dependent manner serving precise cognitive functions ^14^.

Second, while most PAC studies focus on nested gamma oscillations at lower frequencies (35-55 Hz) ^40, 46^, LFPs due to genuine oscillations do appear at high-gamma frequencies, as well. For example, Pernia-Andrade and Jonas ^47^ demonstrated IPSPs in intracellular recordings of DG granule cells coherent with extracellular LFPs in the high-gamma range (76+5 Hz). Cross-frequency coupling of real network oscillations were also shown in this range and even above 100 Hz, e.g. 50-90 and 90-150 Hz ^39^, 100-140 Hz ^22^, 80-150 Hz ^41^, up to 140-180 Hz (related to genuine high frequency oscillations, or HFO ^44, 48, 49^) indicating that these gamma oscillations may provide specific channels of communication separate from low-gamma oscillations ^7, 31^. Although unlikely, the contribution of high-gamma oscillations to the present results cannot be completely excluded. Coupling of gamma power and spike activity in the mixture of LFPs was shown in general ^5, 50-52^ and thus some of the bursting neuronal ensembles might be synchronized in the high-gamma range and co-vary with transients related to action potentials.

## Methods

All surgical and other relevant aspects of the experimental procedure were approved by the Institutional Animal Care and Use Committee

### Data Acquisition

Experiments were performed on male Sprague–Dawley rats, (Charles River Laboratories, MA) treated in accordance with NIH guidelines. All procedures were approved by the Institutional Animal Care and Use Committees of Beth Israel Deaconess Medical Center.

#### Dataset I

Rats were implanted with chronic EEG and EMG electrodes under ketamine-xylazine (70–80 and 10 mg/kg, respectively) anesthesia. Field potentials were recorded using surface screw electrodes placed at equidistant locations along a rostro-caudal axis, over the left frontal, parietal and occipital cortex (AP: 1.0, −2.5, −6.5 mm, Lat: 2.0, 2.5, 3.0 mm, respectively) and with fine wires in the prefrontal cortex (PFC) (AP: 3.2, Lat: 0.2, DV: 5.1 mm, on both sides) and hippocampus (AP: 3.7, Lat: 2.2, DV: 3.5 mm, on the right side). Daily recordings started 7–10 days after surgery. The rats were placed in a recording box in the morning, and cortical and hippocampal EEG, and neck muscle EMG were continuously recorded for 24 h without disturbing the animal ^49, 53^. Theta-intensive periods were selected from awake as well as REM sleep and recordings from the hippocampus and PFC were submitted to analysis (Figure 2). A total of 6 recordings of awake and 6 recordings of REM from 6 animals (n = 6) were used for this experiment. Awake data and REM data were separately analyzed.

#### Dataset II

Rats were prepared for chronic recordings using surgical and recording techniques similar to Dataset I. Hippocampal LFPs were recorded in DG and CA1 regions of the hippocampus using a pair of twisted wires placed above and below the hippocampal fissure (AP: 3.7, Lat: 2.2, DV: 2.5 and 3.5 mm), verified by out-of-phase theta oscillations and later by histology ^54, 55^. LFPs from simultaneous CA1 and DG recordings during awake and REM sleep theta states were submitted to analysis (Figure 2). A total of 13 different recordings from 4 animals (n = 13) were analyzed.

#### Dataset III

Hippocampal LFPs were recorded from rats under urethane anesthesia using a 16-channel linear multicontact electrode with 100 µm separation between contacts. The linear electrode was implanted in the dorso-ventral direction to cover a 1.5–2.5 mm segment across CA1, DG, and hilar regions^12, 18^ (Figure 2). CA1 and DG regions were identified by perforant path evoked potentials. Theta rhythm was elicited by high frequency (100Hz) stimulation of the pontine reticular formation ^12, 18, 54^. A total of 9 different recordings from 5 animals (n = 9) were analyzed.

### Data Analysis

#### Preprocessing

A high pass filter (> 1 Hz) and a notch filter centered at 60 Hz were applied to all datasets. For Granger causality (GC) analysis (Dataset III), LFP data was downsampled from 1k Hz to 200 Hz.

#### Power spectral analysis

Power spectra were obtained using the Welch method of power spectral estimation where the window was 2 sec in duration with 50% overlap.

#### Phase amplitude coupling (PAC)

Each dataset consisted of continuous data segments of varying length. The unit of analysis for PAC were these segments of continuous recordings referred to as recordings. For Datasets I and II, the length of such segments varied between 50 sec and 1000 sec, whereas for Dataset III, the length of recordings varied between 5 sec and 10 sec. The PAC analysis method followed that of Canolty et al. (2006). First, consider two LFP signals. One was filtered in the high-gamma frequency range (65 – 85 Hz), denoted fA, and the other in the theta frequency range (4 – 8 Hz), denoted fp. From the Hilbert transforms of the filtered signals, the amplitude of high-gamma, denoted 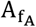, and the phase of theta, denoted Φ_fp_, were extracted, and the complex-valued composite analytic signal 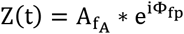 was formed. The modulation index (MI) was derived from the first moment (M_RAW_) of Z(t). M_RAW_ was normalized before it was used as a metric of coupling strength. Surrogate means were created by introducing lag τ between 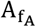 and Φfp such that Z (t, τ) = 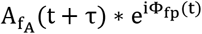. Normalized Z-score was obtained to yield M_NORM_ = (M_RAW_ – μ)/σ where μ was the mean of the surrogate lengths and σ was their standard deviation. M_NORM_ was used as a measure of interregional PAC or ir-PAC. For Local PAC or l-PAC, the filtering was done on the same LFP signal, and the other steps were the same. To compute PAC comodulogram, M_NORM_ was calculated between amplitude of LFP signal ranging from 1 to 100 Hz with 10 Hz bandwidth and 5 Hz step and phase of LFP signal ranging from 1 to 30 Hz with 2 Hz bandwidth and 0.5 Hz step.

#### Granger causality analysis

Parametric GC analysis was performed on Dataset III. Bipolar signals were derived from the theta generators in DG and CA1 respectively and the two bipolar signals were then subjected to AR modeling (see Trongnetrpunya et al., 2015) from which Granger causality spectral estimates were obtained. For each stimulation strength, the data were divided into epochs of 2 seconds in duration. Each epoch was treated as a realization of an underlying stochastic process. Here, bipolar derivation was taken to eliminate common signal and volume conduction, and shown to be a critical step in the proper estimation of Granger causality between two neuronal ensembles ^12^. For Datasets I and II, due to the electrodes applied, bipolar derivation was not possible to obtain. Thus, Granger causality analysis was not attempted on these two datasets.

#### Testing of statistical significance

To test whether the estimated Granger causality or the PAC (l-PAC or ir-PAC) modulation index was significantly greater than 0, we utilized a random permutation approach ^56^. In this approach a baseline null-hypothesis distribution was constructed from which statistical significance threshold was derived. Each continuous data recording was divided into epochs of 2 second in duration. For two time series within the segment the epoch index from one was permuted randomly against that from the other to create a synthetic dataset. Granger causality spectra and PAC modulation index were derived from the synthetic dataset and the appropriate average value was taken. This random permutation procedure was repeated five hundred times to yield the null hypothesis distribution of GC and PAC spectra for the respective statistical significance test. In both the cases, values from the actual dataset were compared against the synthetic data distribution and considered significant only if they exceeded the 95th percentile value of the null hypothesis distribution (p < 0.05).

## Acknowledgement

This work was supported by the National Institute of Mental Health grant R01 MH100820.

## Author Contributions

BN, BK, and MD interpreted the results and wrote the manuscript, BN and MD analyzed the data, PS and BK performed the experiments.

